# Multiple knockout mutants reveal a high redundancy of phytotoxic compounds that determine necrotrophic pathogenesis of *Botrytis cinerea*

**DOI:** 10.1101/2021.08.21.457223

**Authors:** Thomas Leisen, Janina Werner, Patrick Pattar, Edita Ymeri, Frederik Sommer, Michael Schroda, David Scheuring, Matthias Hahn

## Abstract

*Botrytis cinerea* is a major pathogen of more than 1400 plant species. During infection, the kills host cells during infection and spreads through necrotic tissue, which is believed to be supported by induction of programmed plant cell death. To comprehensively evaluate the contributions of most of the currently known plant cell death inducing proteins (CDIPs) and metabolites for necrotrophic infection, an optimized CRISPR/Cas protocol was established which allowed serial marker-free mutagenesis to generate Botrytis mutants lacking up to 12 different CDIPs. Infection analysis revealed a decrease in virulence with increasing numbers of knockouts, and differences in the effects of knockouts on different host plants. The *on planta* secretomes obtained from these mutants revealed substantial remaining necrotic activity after infiltration into leaves. Our study has addressed for the first time the functional redundancy of virulence factors of a fungal pathogen, and demonstrates that *B. cinerea* releases a highly redundant cocktail of proteins and metabolites to achieve necrotrophic infection of a wide variety of host plants.

## Introduction

*Botrytis cinerea* is considered as one of the most important plant pathogenic fungi, causing severe pre- and postharvest losses on fruits, vegetables and other crops worldwide (Elad Y *et al*. 2016). The fungus attacks its host plants preferentially under humid and cool conditions. Before and after invasion into the plant tissue, the hyphae kill the surrounding host cells and spread through the dying tissue, followed by the development of a superficial mycelium which has the typical grey mold appearance and releases a plethora of conidia into the air. Mechanisms that have been proposed to promote necrotrophic infection of *B. cinerea* are the secretion of plant cell death inducing proteins (CDIPs) and cell wall degrading enzymes, the release of phytotoxic metabolites and organic acids, and the acidification of the host tissue (Müller *et al*. 2018; Veloso and van Kan, 2018; Zhu *et al*. 2017). Furthermore, the fungus can suppress host defense gene expression by the release of small interfering RNAs (Weiberg *et al*. 2013), and it is able to detoxify plant defence compounds such as camalexin and tomatine via efflux transporters or by enzymatic modification (Stefanato *et al*. 2009; You and van Kan, 2021).

How host cell death is induced is not fully understood, but there is evidence that necrotrophic fungi actively trigger the hypersensitive response (HR), a plant-specific type of programmed cell death linked to strong defence reactions including an oxidative burst (Govrin and Levine, 2000; Veloso and van Kan, 2018). Several secreted compounds have been described as virulence factors. *B. cinerea* releases two major phytotoxic metabolites, the sesquiterpenoid botrydial and the polyketide botcinin, which have been shown to be together required for full virulence (Dalmais *et al*. 2011; Pinedo *et al*. 2008). Plant cell wall degrading enzymes (CWDE) are required for tissue mazeration by necrotrophic pathogens, but because of their redundancy, the contributions of individual members are difficult to determine. *B. cinerea* mutants lacking either of the two major endopolygalacturonases, PG1 and PG2, showed impaired lesion formation (Have *et al*. 1998; Kars *et al*. 2005). An endo-arabinanase (BcAra1) was found to be required for full infection of Arabidopsis but dispensable for infection of tobacco (Nafisi *et al*. 2014). Further, a cellobiohydrolase and a β-endoglucanase were reported to contribute to plant infection (Li *et al*. 2020). Several CWDEs of *B. cinerea* are CDIPs, inducing necrosis of different plant tissues. Necrotic activity was found to be independent of enzymatic activity for two xylanases, Xyn11A and Xyl1, and the xyloglucanase XYG1. For Xyn11A and Xyl1, peptides of 25 and 26 amino acids, respectively, were identified that induced cell death (Frías *et al*. 2019; Yang *et al*. 2018), and for XYG1 two exposed loops of the folded protein were identified as being essential to induce cell death (Zhu *et al*. 2017). Mutants lacking Xyn11A and Xyl1 showed impaired infection (Noda *et al*. 2010; Yang *et al*. 2018). *B. cinerea* also secretes CDIPs without known enzymatic activity. Nep1 and Nep2, which belong to a large family of plant necrosis and ethylene inducing proteins in fungi, oomycetes and bacteria, induce pores in membranes of dicotyledonous plants (Seidl and van den Ackerveken, 2019). *B. cinerea* mutants lacking either Nep1 or Nep2 showed normal virulence (Cuesta Arenas *et al*. 2010). The cerato-platanin Spl1, was found to be required for full infection, whereas elimination of IEB1 had no effect on virulence (Frías *et al*. 2011; Frías *et al*. 2016). The recently identified CDIP Hip1 was found to require its tertiary structure for phytotoxic activity. While knockout mutants showed normal infection, strains overexpressing Hip1 showed revealed slightly increased virulence compared to the wild type (WT) strains (Frías *et al*. 2016; Jeblick *et al*. 2020).

Similar to infection by *B. cinerea*, treatment of leaf tissues with individual CDIPs usually results in HR-like programmed cell death (Frías *et al*. 2011; Frías *et al*. 2013; Frías *et al*. 2016; Yang *et al*. 2018; Zhu *et al*. 2017). CDIPs oft act as pathogen associated molecular patterns (PAMPs) that are recognized by pattern recognition receptors (PRRs) in the plant membrane, leading to the so-called PAMP-triggered immunity (PTI) (Thomma *et al*. 2011). A PAMP-like behavior of CDIPs was indicated by the observation that their activity was dependent on the presence of the PRR coreceptors BAK1 and/or SOBIR1 in the treated plants (Franco-Orozco *et al*. 2017; Frías *et al*. 2011; Yang *et al*. 2018; Zhu *et al*. 2017), and PRRs for Nep1/Nep2, Xyn11A and PG1/PG2 have already been identified (Albert *et al*. 2015; Ron and Avni, 2004; Zhang *et al*. 2014). Recently, a novel *B. cinerea* CDIP was discovered that is translocated into plant cells, but it does not seem to contribute to infection (Bi *et al*. 2021).

Based on the data summarized above, it is evident that *B. cinerea* secretes a mixture of phytotoxic compounds to kill host cells, and triggering of HR seems to play an essential role in this process. Because elimination of single CDIPs or phytotoxins has been shown to have either no or only limited effects on pathogenesis, a comprehensive approach is required for understanding how and to what extent CDIPs contribute to necrotrophic pathogenesis. The goal of this study was to create single and multiple mutants of genes for most of the currently known CDIPs and phytotoxic metabolites, in the same genetic background and one laboratory, and to evaluate their contribution to the infection process of *B. cinerea*. By applying an improved version of a recently developed CRISPR/Cas9-based method for marker-free genome editing (Leisen *et al*. 2020), we generated multiple mutants lacking up to 12 CDIPs and phytotoxins. These mutants were unaffected in their growth and differentiation *in vitro*, but showed significantly impaired virulence compared to WT on different host tissues. These data highlight the role and the complexity of the toxic secretome for necrotrophic infection of *B. cinerea*.

## Results

### Generation of *B. cinerea* single and multiple CDIP mutants

Before the CRISPR/Cas protocol for marker-free mutagenesis was available (Leisen *et al*. 2020), different selection markers were used to generate single and up to quadruple mutants by standard mutagenesis, using knockout constructs integrated into the target genes by homologous recombination. In addition to the established markers conferring resistance to hygromycin (HygR), nourseothricin (NatR) and fenhexamid (FenR), the anilinopyrimidine fungicide cyprodinil (Cyp)(Heye *et al*. 1994) was developed as a new marker for selection in *B. cinerea*. The mode of Cyp resistance (CypR) has been uncovered by a functional genomics approach (Mosbach *et al*. 2017). Many CypR *B. cinerea* field strains contain a mutation leading to an L412F exchange of a mitochondrial NADPH kinase encoded by *Bcpos5*. A CypR selection marker was generated by integrating *Bcpos5*^*L412F*^ into a constitutive expression cassette (Fig. 1A). Functionality of the CypR marker was confirmed by targeted mutagenesis of several genes, which yielded robust numbers of transformants, with a low fraction of spontaneous CypR mutants (Fig. 1B,C).

**Figure 1:**
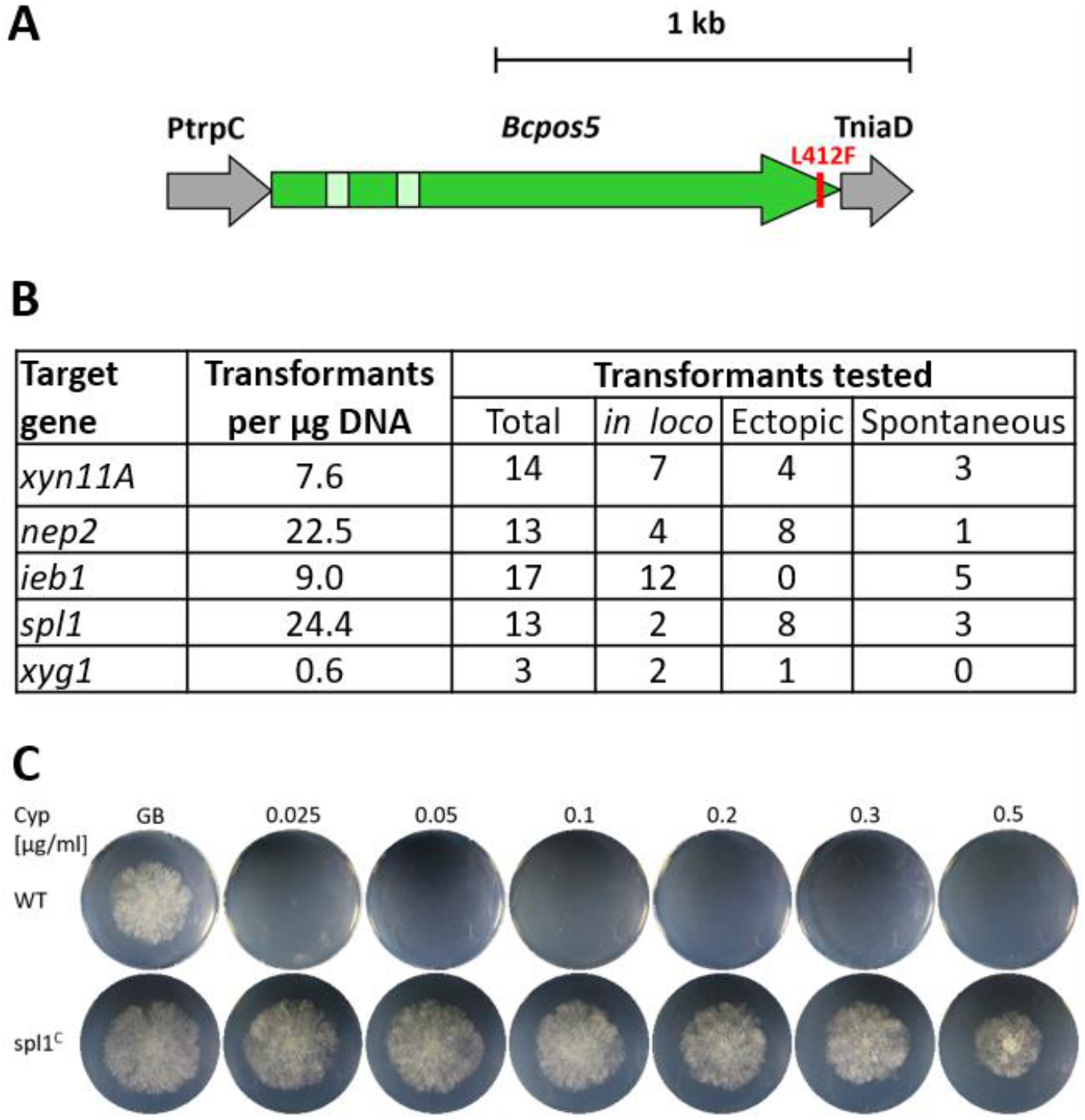
Establishment of cyprodinil resistance (CypR) as a novel selection marker for *B. cinerea*. A: CypR cassette containing a 344 bp promoter fragment of *Aspergillus nidulans trpC*, and a 146 bp terminator fragment of *B. cinerea niaD*. B: Transformation efficiency with the CypR cassette and characterization of the CypR transformants. *in loco*: Correct knockout mutants (replacement of WT DNA by CypR cassette). Ectopic: Detection of CypR cassette outside of target gene. Spontaneous: FenR ‘transformants’ lacking the CypR cassette. C: Growth of a CypR resistant transformant (spl^C^), and the sensitive *B. cinerea* WT strain on agar plates with GB5 minimal medium containing different Cyp concentrations.

The genes analysed by mutagenesis include most of the currently known *B. cinerea* CDIPs and phytotoxins (Table 1). Single knockout mutants were generated for the previously characterized genes *xyn11A, spl1, xyg1, ieb1*, and *nep2*. Furthermore, we constructed mutants of *gs1* encoding a putative glucoamylase previously reported as a CDIP (Zhang *et al*. 2015), and of *plp1* (PAMP like protein) encoding the homolog of a CDIP of the apple pathogen *Valsa mali* named VmE02 (Nie *et al*. 2019). All mutants showed normal vegetative growth and differentiation *in vitro*. When infection experiments were performed with tomato and *Phaseolus* bean leaves and apple fruits, no significant differences in virulence compared to WT were observed (Fig. S1). These results demonstrated that none of the deleted genes alone play a major role for infection on any of the tested plants. These results are inconsistent with previous studies, which reported reduced virulence of *B. cinerea xyn11A*, spl1 *and xyl1* mutants (Brito *et al*. 2006; Frías *et al*. 2011; Noda *et al*. 2010). Next, a quadruple mutant (4x^R^: *xyn11A spl1 nep1 nep2*) was generated by using four different resistance markers, including the newly established CypR marker. As shown below, the 4x^R^ mutant was weakly impaired in virulence, besides a minor growth retardation. This would be consistent with weak phenotypes of the single mutants that are too small to be detected in our infection assays.

**Table 1:** Cell death-inducing (CDI) metabolites and CDIPs of *Botrytis cinerea*

|  | k.o. effects | *in planta* expression<sup>1</sup> | PRR | Evidence for PRR coreceptors | References |
| --- | --- | --- | --- | --- | --- |
| **CDI metabolites** |  |  |  |  |  |
| Botrydial | yes<sup>2</sup> | ++ | no | --- | Dalmais *et al.* 2011 |
| Botcinic acid |  | ++ | no | --- |  |
| **CDIPs (non-enzymatic)** |  |  |  |  |  |
| Nep1 | no | + / +++ | RLP23 | BAK1, SOBIR1 | Cuesta Arenas *et al.* 2010; Albert *et al.* 2015; Ono *et al.* 2020 |
| Nep2 | no | ++ | RLP23 | BAK1, SOBIR1 |  |
| Spl1 (cerato-platanin) | yes | +++ | yes<sup>5</sup> | BAK1 | Frías *et al.* 2011 |
| IEB1 | no | +++ | yes<sup>5</sup> | no | González *et al.* 2017 |
| Hip1 | no | +++ | yes<sup>5</sup> | no | Jeblick *et al.* 2020 |
| PIP1 | no | + | REO2 | BAK1, SOBIR1 | Nie *et al.* 2019, 2021 |
| **CDIPs: Enzymes<sup>3</sup>** |  |  |  |  |  |
| Xylanase Xyn11A | yes | ++ / +++ | LeEIX2 | no | Noda *et al.* 2010; Ron and Avni, 2004 |
| Xyloglucanase Xyg1 | no | ++ | yes | BAK1, SOBIR1 | Zhu *et al.* 2017 |
| Xylanase Xyl1 | yes | + | yes | BAK1, SOBIR1 | Yang *et al.* 2018 |
| Glucoamylase Gs1 | n.a. | ++ | yes | no | Zhang *et al.* 2015 |
| Polygalacturonase PG1 | yes | +++ | RBPG1 | BAK1, SOBIR1 | Have *et al.* 1998; Kars *et al.* 2005; Zhang |
| Polygalacturonase PG2 | yes | +++ | RBPG1 | BAK1, SOBIR1 | *et al.* 2014 |

<sup>1</sup> Based on reads per kilobase million values from RNA sequencing data from infected tomato leaves (Müller *et al.* 2018); Jan van Kan, unpublished); +++: RPKM >1000; ++: RPKM 100-1000; +: RPKM <100.
<sup>2</sup> No effects of single k.o., reduced virulence of double k.o.
<sup>3</sup> CDI activity independent of enzyme activity (except for PG1, PG2)

### Development of a CRISPR/Cas method for rapid generation of homokaryotic, multiple knockout mutants

We have recently described a powerful CRISPR/Cas-based method for *B. cinerea* gene editing without introducing resistance markers into the transformants. It is based on cotransformation of Cas9-sgRNA RNPs with an unstable telomere vector into protoplasts, which allows transient selection of transformants containing the desired editing events (Leisen *et al*. 2020). For improved serial mutagenesis, the protocol was modified to generate deletions by two RNPs targeting one gene, without addition of a repair template, which results in excision of the sequence between the cleavage sites by non-homologous end joining (NHEJ). When the protocol was tested for knockout of *spl1*, several thousand FenR transformants were obtained, and 68% of them verified by PCR analysis as being edited with the correct deletion. For serial mutagenesis, two genes were targeted simultaneously by using four RNPs in each transformation. These experiments usually yielded high editing rates, resulting in the isolation of mutants containing the expected single and double deletions (Table 2). Unexpectedly, a large fraction of the primary transformants appeared to be homokaryotic, and complete loss of the deleted DNA in these transformants was confirmed after a single spore isolation step. This represents a great advancement over traditional transformation methods for *B. cinerea*, which usually resulted in heterokaryotic transformants that had to be purified by several rounds of single spore isolation before homokaryosis was achieved (Hahn and Scalliet, 2021).

**Table 2:** Serial deletion of *B. cinerea* genes encoding CDIPs and phytotoxic metabolites.

| Transf. Round | Strain (mutant) used for transformation |  | Genes targeted for k.o. | Characterization of transformants |  |  |  |  |  |  |
| --- | --- | --- | --- | --- | --- | --- | --- | --- | --- | --- |
| | Name | Genotype | | Total | Tested | Deletion 1* | Homo-karyons | Deletion 2* | Homo-karyons | $\Delta 1 + \Delta 2^*$ |
| 1 | WT | WT | *spl1* | >1000 | 22 | 15 (68%) | 6 | n.a. | n.a. | n.a. |
| 2 | spl1<sup>&</sup> | *spl1* | *nep1, nep2* | 157 | 26 | 6 (23%) | 4 | 7 (27%) | 5 | 2 (8%) |
| 3 | 3x<sup>&</sup> | *spl1 nep1 nep2* | *xyn11A, ieb1* | >1000 | 104 | 13 (13%) | 10 | 6 (6%) | 4 | 0 (0%) |
| 4 | 4x<sup>&</sup> | *spl1 nep1 nep2 xyn11A* | *hip1, xyg1* | >5000 | 70 | 57 (81%) | 50 | 54 (77%) | 49 | 40 (57%) |
| 5 | 6x | *spl1 nep1 nep2 xyn11A hip1 xyg1* | *plp1, ieb1* | >1000 | 46 | 20 (43%) | 17 | 10 (22%) | 8 | 4 (9%) |
| 6 | 8x | *spl1 nep1 nep2 xyn11A hip1 xyg1 plp1 ieb1* | *xyl1, gs1* | >1000 | 64 | 31 (48%) | n.a. | 8 (13%) | n.a. | 8 (13%) |
| 7a | 10x | *spl1 nep1 nep2 xyn11A hip1 xyg1 plp1 ieb1 xyl1 gs1* | *pg1, pg2* | 102 | 56 | 5 (9%) | 5 | 1(2%) | 1 | 1 (2%) |
| 7b |  |  | *bot2, boa6* | >1000 | 105 | 3 (3%) | 2 | 26 (25%) | n.a. | 3 (3%) |
|  | Name | Genotype of mutants derived from strain 10x transformed with *pg1, pg2* (11x, 12xpg) or *bot2, boa6* (12xbb) |  |  |  |  |  |  |  |  |
|  | 11x | *spl1 nep1 nep2 xyn11A hip1 xyg1 plp1 ieb1 xyl1 gs1 pg1* |  |  |  |  |  |  |  |  |
|  | 12xpg | *spl1 nep1 nep2 xyn11A hip1 xyg1 plp1 ieb1 xyl1 gs1 pg1 pg2* |  |  |  |  |  |  |  |  |
|  | 12xbb | *spl1 nep1 nep2 xyn11A hip1 xyg1 plp1 ieb1 xyl1 gs1 bot2 boa6* |  |  |  |  |  |  |  |  |

\*Number and percentage of transformants with confirmed deletions in gene 1 ( $\Delta 1$ ), gene 2 ( $\Delta 2$ ), or both genes.
<sup>&</sup> Mutants were not phenotypically characterized.

The improved transformation protocol was used for serial inactivation of up to 12 genes encoding CDIPs and key enzymes for biosynthesis of the phytotoxins, botrydial and botcinin. The expected gene deletions, and the absence of remaining WT DNA in the deleted regions were verified by PCR (Fig. S2). Sequencing revealed in most cases precise (±2 bp) excisions as predicted from the RNP-directed DNA cleavage sites, except for the Δ12xpg mutant in which a 3 kb larger deletion than expected had occurred in *pg2* (Table S1).

### Phenotypic analysis of multiple mutants lacking up to 12 CDIPs and phytotoxins

All mutants displayed growth and sporulation similar to WT (Fig. 2A, B). Sclerotia formation, which is induced by cultivation in complete darkness, was also unaffected (Fig. 2C). When incubated on glass surfaces for two days, WT germlings form large aggregates of appressoria-like structures, so-called infection cushions. They are believed to represent alternative infection structures to simple appressoria (Choquer *et al*. 2021). Infection cushions with similar morphology were formed by the WT and the 12xpg and 12xbb mutants (Fig. 2D). When inoculated onto killed onion epidermal layers, conidia WT and mutants showed a similar infection behaviour, by forming short germ tubes following by penetration into host cells and formation of thick intracellular hyphae (Fig. 2E). These data confirmed that none of the secreted proteins are involved in vegetative growth, reproduction and pathogenic differentiation on artificial surfaces and killed host tissue.

**Figure 2:**
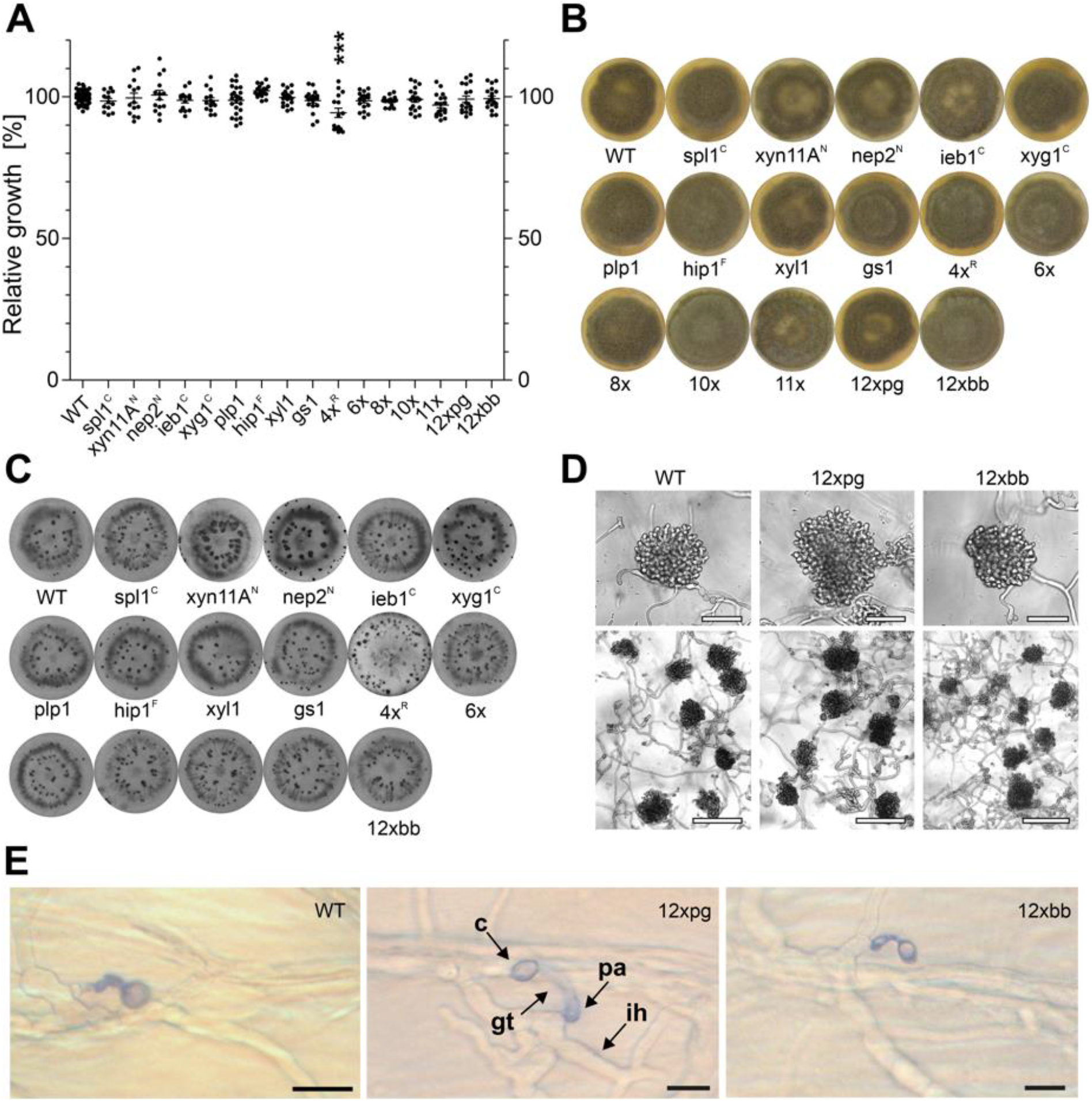
Growth and differentiation of *B. cinerea* B05.10 (WT) and mutants generated in this study. A: Relative radial growth on GB5 minimal agar medium with 25 mM glucose (3 days). The p values by one way ANOVA followed by Dunnett’s multiple comparisons post-hoc test are indicated; ***: p < 0.001. B: ME plates incubated for 10 days under permanent light to induce conidia formation. C: ME plates incubated for 14 days in darkness to induce sclerotia formation. Mutants marked with superscript letters were generated with resistance markers. D: Infection cushions formed on glass slides after 48 h. Upper scale bars: 50 µm; lower scale bars: 150 µm. E: Penetration of onion epidermis cells by germinated conidia of WT, 12xpg and 12xbb mutants. Superfical structures (conidium, germ tube and appressorium) are stained with trypan blue, whereas intracellular hyphae remain unstained. Scale bars: 10 µm.

To test the effects of multiple knockouts on infection, leaves of *Phaseolus* bean, tomato and maize, and apple fruits were inoculated, and necrosis formation quantified after 48 to 96 h (Fig. 3). On all tested tissues, infection efficiency of the mutants decreased with increasing number of deleted genes. On bean leaves and apple fruit, the mutants revealed a stronger reduction in virulence than on tomato and maize leaves. For example, the 10x mutant formed lesions which were only ca. 60% in size of WT lesions on bean leaves and apple, but similar or only slightly smaller lesions on tomato and maize leaves. Compared to the 10x mutant, 11x, 12xpg and 12xbb mutants showed further reductions of lesion sizes, except on maize leaves. These comparisons allowed to assign contributions to virulence for *pg1* (10x vs. 11x), *pg1* plus *pg2* (10x vs. 12xpg), encoding endopolygalacturonases, and *bot2* plus *boa6* encoding key biosynthesis enzymes for botrydial and botcinin (Fig. 3). The effects of *pg1* and *pg2* were confirmed by the reduced virulence of a *pg1 pg2* double mutant mutant, generated by classical mutagenesis with selection markers, on all tested tissues (Fig. S3), in accordance to previous results (Have *et al*. 1998; Kars *et al*. 2005). The effects of *bot2* and *boa6* knockouts were most evident by the low virulence of the 12xbb mutant on apple. This was confirmed by the infection phenotype of a previously generated *bot2 boa6* double mutant (Leisen *et al*. 2020), which showed considerable reduction in lesion formation on apples (Fig. S4). These results are consistent with published data (Dalmais *et al*., 2011). Despite their reduced virulence, the 12xpg and 12xbb mutants were eventually able to sporulate on infected bean and tomato leaves, similar to WT (Fig. S5).

**Figure 3:**
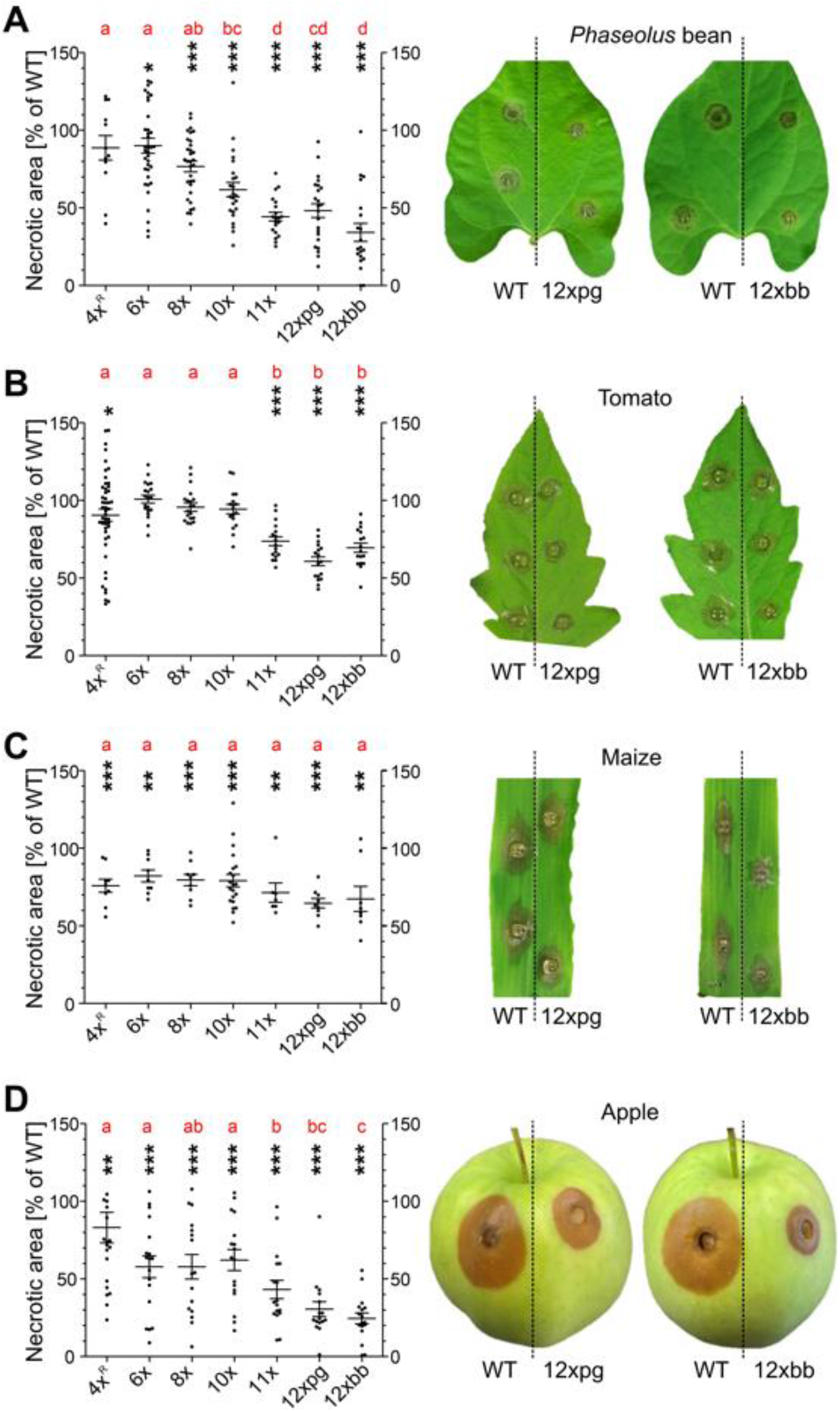
Infection tests of multiple CDIP/ phytotoxin mutants. A: Attached *Phaseolus* bean leaves (48 h). B: Detached tomato leaves (48 h). C: Detached maize leaves (72 h). D: Apple fruits (96 h). The p values by one-sample t test to a hypothetical value of 100% (WT) are indicated. * p value < 0.05; ** p value < 0.01; *** p value < 0.001. Results of Tukey’s multiple comparison test are displayed with compact letter display. The pictures show lesions caused by 12xbb and 12xpg mutants in comparison to WT.

### Microscopic analysis of infection of WT and multiple mutants

The infection process of *B. cinerea* can be divided into penetration, primary lesion formation, lesion expansion and sporulation. To investigate whether the reduced virulence of the 12x mutants was related to defects in early stages of differentiation and infection, microscopic studies were performed. On *Phaseolus* leaves, host cell killing by both mutants was substantially reduced after 24 h (Fig. 4). These data show that the CDIPs deleted in these mutants are involved in the early stages of lesion formation on unwounded leaf tissue.

**Figure 4:**
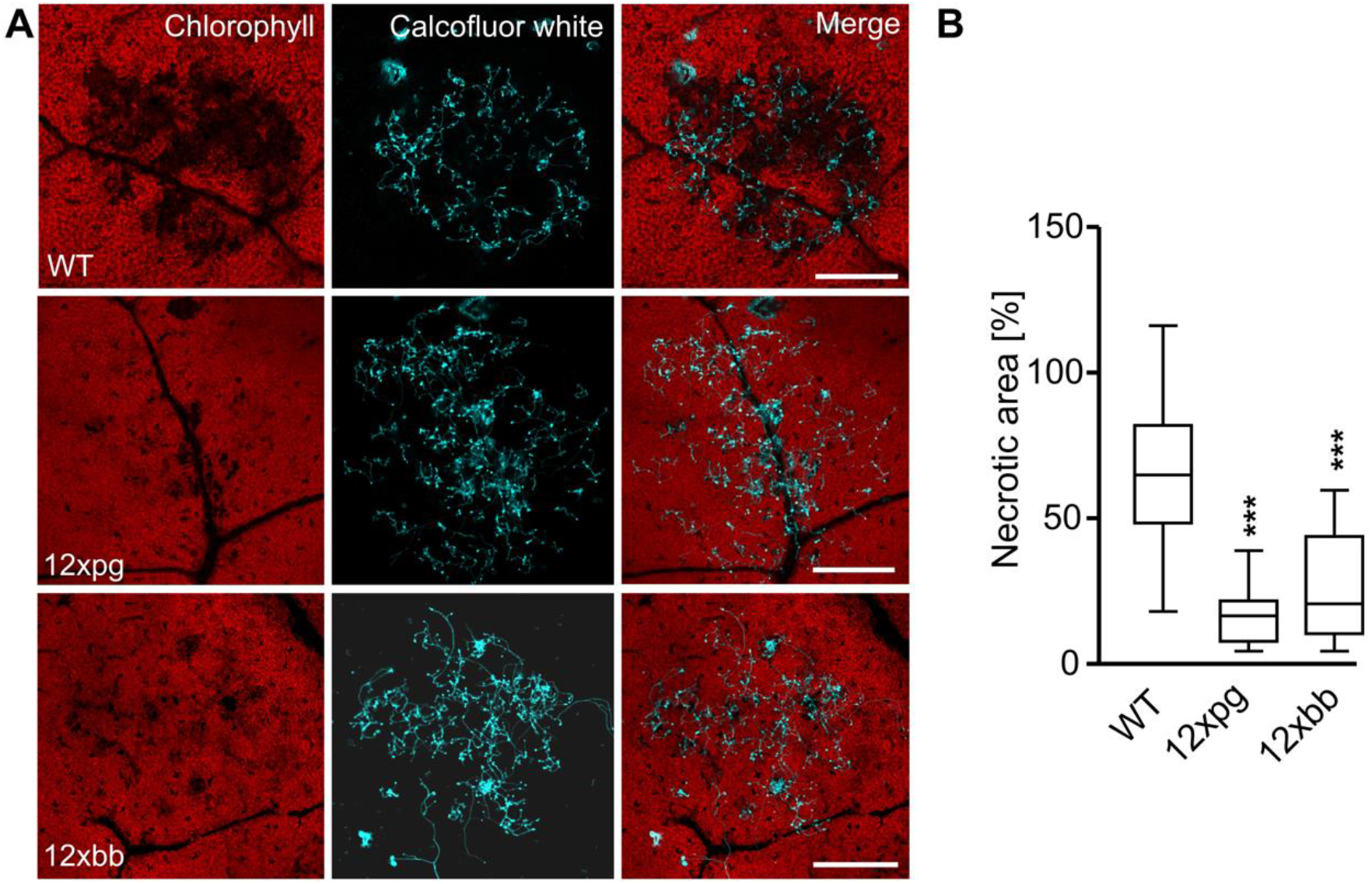
Microscopic analysis of early stage of infection of *B. cinerea* WT and mutants. Fungal hyphae were stained with calcofluor white, host cell death is visible by loss of autofluorescence. A: Host cell killing on *Phaseolus* leaves (24 h). Scale bars: 500 µm. B: Quantification of host necrosis. One-way ANOVA followed by Dunnett’s multiple comparison post hoc test ***p<0.001.

### Analysis of the secretomes of WT and multiple mutants

To verify the loss of proteins encoded by the deleted genes in the mutants, an proteomic analysis of *on planta* produced secretomes was performed according to (Müller *et al*. 2018). Out of the 12 CDIPs analysed, nine could be detected in the WT secretome, consistent with previous studies (Zhu *et al*. 2017; Müller *et al*. 2018). In all mutants, proteins were missing when the respective gene had been deleted (Table S2). Since most of the deleted CDIPs are highly expressed in the WT, we checked whether their loss was compensated by overexpression of other proteins in the secretomes of the 10x, 11x, 12xpg and 12xbb mutants. However, analysis of the proteome data using the Perseus bioinformatic platform (Tyanova and Cox, 2018) did not reveal evidences for differential protein abundance in the WT and mutant secretomes.

The *on planta* secretomes of *B. cinerea* are highly phytotoxic when infiltrated into leaves (Jeblick *et al*. 2020; Zhu *et al*. 2017). Similar toxicity was observed for the secretomes of WT and up to 8x mutants. The secretomes of 10x, 12xpg and 12xbb mutants showed reduced cell death inducing activity. Compared to 10x, the secretomes of 12xpg and 12xbb showed decreased activity on *V. faba* and *N. benthamiana*, respectively, indicating differential effects of the loss of PG1/PG2 (in 12xpg) and botrydial/botcinin (in 12xbb) on the different plant species (Fig. 5A-D). As the 10x and the 12xbb mutant differed in their ability to synthesize botrydial and botcinin, we investigated the contribution of the two phytotoxins. Secretomes collected from the two mutants were fractionated by ultrafiltration through a membrane with 10 kDa molecular weight cut off. No protein could be detected in the filtrates, and only the filtrate of the 10x but not the 12x mutant caused necrosis, which could be attributed the presence and absence of botrydial and botcinin, respectively (Fig. 5E). Heating of the 10x mutant filtrate to 95°C for 20 min did not signficantly reduce its phytotoxic activity (Fig. 5F).

**Figure 5:**
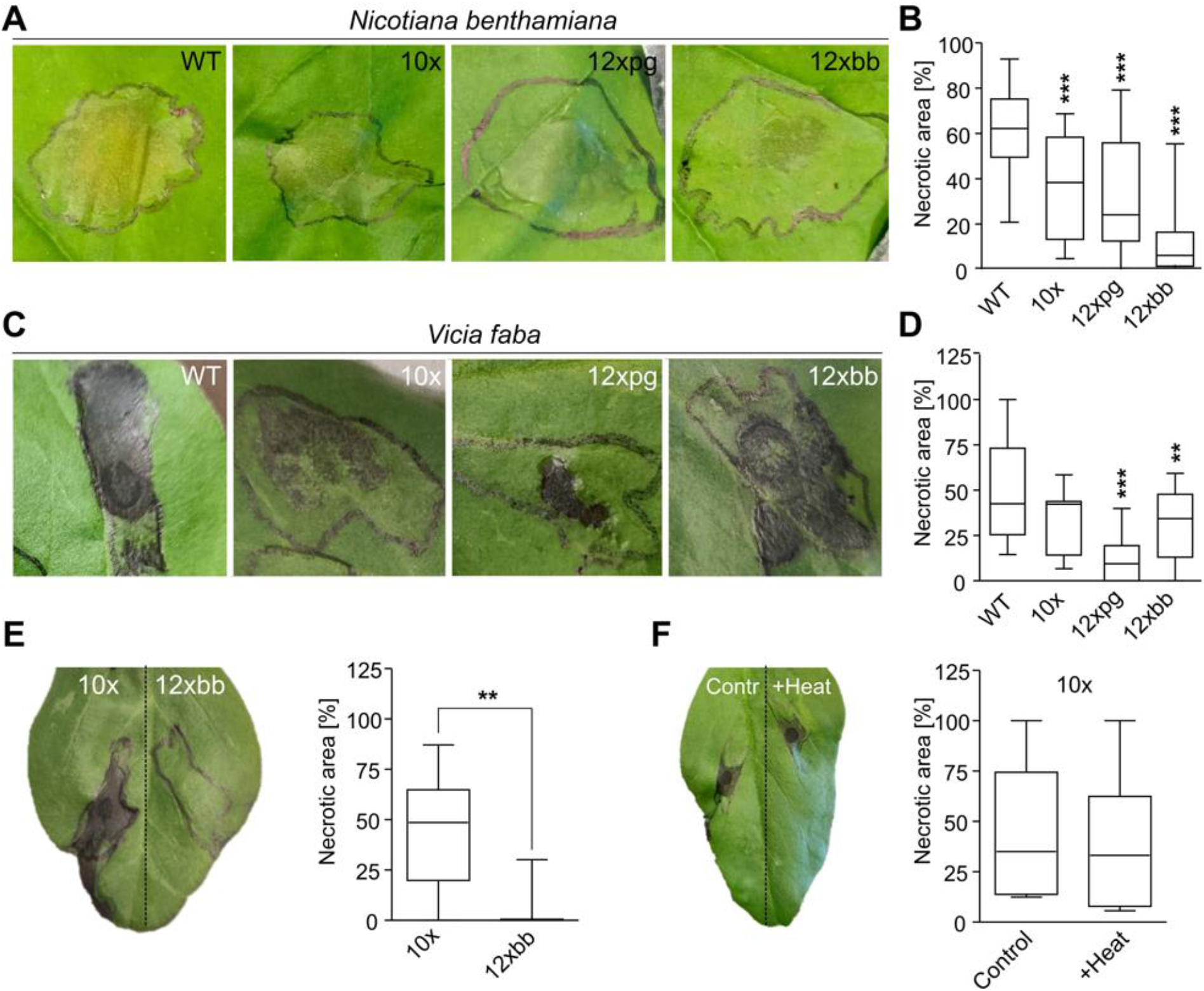
Cell death inducing activity of *B. cinerea on planta* secretomes. A-D: Necrotic lesions caused by WT and mutant secretomes (2 µg ml^-1^) in infiltrated tobacco (A,B) or faba bean (C,D) leaves. A, C: Lesions formed in infiltrated leaf areas. B, D: Quantitative evaluation of CDI activity of WT and mutants’ secretomes. Values are the means of at least three experiments and two or three leaves per experiment. One-way ANOVA followed by Dunnett’s multiple comparison post hoc test; ***p<0.01; ***p<0.001 (n≥15 for tobacco, n≥8 for faba bean). E: Cell death inducing activity on faba bean leaves of non-proteinaceous (<10kDa) secretome fractions of a 10x mutant and a 12xbb mutant unable to synthesize botrydial and botcinin. F: Effects of heating (95°C for 20 min) on CDI activity of the non-proteinaceous fraction of the 10x mutant. E, F: Student t-test; **p<0.01 (n=6).

## Discussion

For the generation of multiple mutants with standard mutagenesis techniques, the availability of selection markers is quickly becoming limiting. With a mutated version of *Bcpos5* conferring resistance to the fungicide cyprodinil, we have established a new selection marker which works with similar efficiency as the established markers HygR, FenR and NatR. The advantage of the fungicide resistance markers FenR and CypR is their low cost, because selection can be applied with commercial fungicide formulations. Using the four available selection markers, we have constructed a 4x^R^ mutant (*spl1 xyn11A nep1 nep2*). Compared to WT and the marker-free mutants generated with CRISPR/Cas, this mutant showed a slight growth retardation. Whether this is due to the constitutive expression of the resistance genes is unclear, but highlights a disadvantage of their use for mutant generation. Based on a recently developed CRISPR/Cas method, we have further improved and simplified the protocol for serial introduction of marker-free gene deletions into *B. cinerea*. A highly favorable result was the high proportion of homokaryotic mutants among the primary transformants. *B. cinerea* protoplasts are generated from germlings containing several nuclei, therefore transformants obtained by standard mutagenesis had to be purified via several rounds of single spore isolations (Noda *et al*. 2007). The reason for the rapid homokaryotization is unclear: We assume that either the RNP complexes are able to edit all nuclei in a protoplast, or that only the nucleus that was edited was able to divide in the transformant. The current protocol allows the generation, verification and purification of multiple *B. cinerea* mutants within three to four weeks. We are not aware of reports in which mutants with a similar number of knockouts have been generated in filamentous fungi until now, besides an eight-fold deletion mutant constructed with a non-CRISPR marker replacement approach in *Aspergillus fumigatus* (Hartmann *et al*. 2011). This and other CRISPR/Cas-based strategies now allow to investigate genes and protein families with redundant functions in most filamentous fungi, except for obligate biotrophs for which any stable transformation remains a great challenge (Martínez-Cruz *et al*. 2017).

We have evaluated the role of 12 CDIPs and two phytoxica metabolites by the construction of single and multiple mutants. In agreement with previous reports, mutants in *nep1, nep2, xyg1* and *hip1* showed normal virulence (Cuesta Arenas *et al*. 2010; Jeblick *et al*. 2020; Zhu *et al*. 2017), as did *gs1* and *plp1* mutants, which have not been described previously. An unexpected result was the absence of significant virulence defects in mutants lacking Xyn11A, Spl1 or Xyl1. These CDIPs have previously been described as virulence factors, based on the analysis of mutants which were also generated in *B. cinerea* strain B05.10 (Brito *et al*. 2006; Frías *et al*. 2011; Yang *et al*. 2018). The discrepancy of these results with our data is difficult to explain, even more as complementation of the published mutants confirmed their reversion to WT phenotypes. A minor role for virulence of *xyn11A* and *spl1* was indicated by the phenotype of the 4x^R^ mutant. Of the six polygalacturonases encoded in the *B. cinerea* genome, PG1 and PG2 are most highly expressed on transcriptional and proteomic levels early during infection of different tissues (Blanco-Ulate *et al*. 2014; Have *et al*. 2001; Müller *et al*. 2018), and both enzymes have been reported to be required for full virulence (Have *et al*. 1998; Kars *et al*. 2005). Accordingly, the virulence defects of the 11x mutant devoid of *pg1* and the 12x mutant lacking *pg1* and *pg2* were also significantly stronger than that of the 10x mutant, which confirmed the effects of the loss of one or both PG isoforms. Further support came from the analysis of an independently constructed *pg1 pg2* mutant, which showed delayed infection on all tested tissues. Considering the redundancy of genes encoding pectin degrading enzymes in the genome of *B. cinerea* (Amselem *et al*. 2011), it is doubtful whether deletion of the remaining endo-PGs would lead to significant further reduction of virulence. Our work also confirms a previous study about the role of the phytoxic metabolites botrydial and botcinin for infection of *B. cinerea* (Dalmais *et al*., 2011). Here, the 12xbb mutant unable to synthesize the two toxins were significantly less virulent than its 10x mutant parent, on all host tissues tested except on maize leaves. This was confirmed by further phenotypic analysis of a *bot2 boa6* double mutant generated previously (Leisen *et al*. 2020). Beyond its phytotoxicity, botrydial has been shown to have antibacterial properties (Vignatti *et al*. 2020). Recently, we have discovered a group of *B. cinerea* field strains which lack the complete botcinin biosynthesis cluster. These strains are less virulent than other *B. cinerea* strains on tomato leaves but not on other host tissues (Plesken *et al*. 2021). These data confirm that botrydial and botcinin together play a significant role for *B. cinerea* pathogenesis, but their mode of action is unknown, and their individual contributions to the infection process remain unclear.

The effects of multiple gene knockouts were variable between host tissues: While smaller effects of multiple gene knockouts on virulence were observed on tomato and maize leaves (lesion sizes >60% of WT), multiple knockouts were considerably less virulent on bean leaves and apple fruit (lesion sizes down to 30% of WT). CDIPs are known to have low plant species specificity, in contrast to many effector proteins from biotrophic and hemibiotrophic fungi. Nevertheless, differences in sensitivity to CDIPs could be due to the presence of different sets of matching receptors or targets in different plant species, or different effects of their activation on plant cell death and defence. PRRs of *B. cinerea* CDIPs belong to the group of receptor like proteins (LRR-RLPs), which have a plant genus- or subgenus-specific distribution (Albert *et al*., 2020). For example, the receptor of PG1/PG2, RBPG1 has been identified in some but not all accessions of Arabidopsis (Zhang *et al*. 2014). No evidence for similar PG receptors exist in tobacco and broad bean leaves, which respond with necrosis only after treatment with enzymatically active but not inactive PGs (Kars *et al*. 2005). Similarly, Hip1 was found to be highly toxic to tobacco but only weakly active in Arabidopsis (Jeblick *et al*. 2021). Furthermore, the membrane-directed toxicity of necrosis and ethylene inducing proteins including Nep1 and Nep2 is known to be restricted to dicots (Lenarčič *et al*. 2017).

The virulence defects observed for the multiple knockout mutants were less pronounced than expected from the high expression levels of most CDIPs and phytotoxins, and from the virulence phenotypes reported for single knockout mutants in previous studies. Because of the lack of virulence defects in the single mutants and only small incremental differences in virulence between the multiple mutants order, it was difficult to estimate the role of single CDIPs, and it remained unclear if all of them contribute to infection. In case of additive and redundant effects, sequential knockouts would result in a stepwise decrease in virulence. This could uncover virulence effects that are too small to be detected in single mutants, for example in *spl1, xyn11A, nep1* and *nep2* in the 4x^R^ mutant. In case of synergism between CDIPs, a stronger decrease in virulence of a multiple mutant than expected from the contributions of the individual gene knockouts would be observed. Synergism has been shown for botrydial and botcinin, since only the double mutant but not the single mutants revealed significant effects on virulence (Dalmais *et al*. 2011). A third possibility, referred to as ‘overkill’, assumes that the total CDI activity of the WT exceeds the requirement for pathogenesis under the chosen infection conditions, and predicts that effects on virulence become evident only when the remaining CDI activity falls below a certain threshold. In this case, deletion of few CDIPs would cause no effects, but the decrease in virulence would be stronger when more CDIPs are deleted. Apart from the botrydial/ botcinin synergism, the moderate but significant virulence phenotypes of the multiple mutants argue for predominantly additive effects. An overkill mechanism still seems possible, and could be revealed if further CDIP knockouts would show increasing effects.

Microscopic analysis revealed a delay of the 12xbb and 12xpg mutants in the early stages of infection. These data demonstrate a role of one or several of the deleted CDIPs in early stages of host attack, in agreement with observations made with *B. cinerea* strains overexpressing *xyg1* which showed evidence for accelerated infection (Zhu *et al*. 2017). However, the markedly reduced infection of all multiple mutants on apple fruits, which are inoculated via wounds, show that several of the deleted CDIPs are also involved in lesion expansion.

The *on planta* secretome of *B. cinerea* is highly phytotoxic, causing cell death in leaf tissue even after five-to ten-fold dilution, down to concentrations of 1 µg ml^-1^. The secretomes of the 10x and 12x mutants were less toxic compared to WT, but retained substantial phytotoxic activity which could be attributed mostly to the protein fraction. Comparison of the 10x and 12xbb mutant secretomes allowed to assign the remaining, heat-stable phytotoxic activity in the low molecular weight (non-proteinaceous) fraction to botrydial and botcinin. Since this fraction was almost nontoxic in the 12xbb mutant, we conclude that probably no other phytotoxic metabolites are secreted in significant amounts during infection of *B. cinerea*. Therefore, the remaining phytotoxic activity in the 12x mutants is due to further, as yet uncharacterized CDIPs. These include Crh1, a newly described CDIP that has been shown to be translocated via infection cushions into host cells (Bi *et al*. 2021). Several CDIPs have been recently described in two molds related to *B. cinerea, Monilinia fructigena* (Vilanova *et al*. 2021) and *Sclerotinia sclerotiorum* (Seifbarghi *et al*. 2020), and homologs for them also exist in *B. cinerea*.

Taken together, our data demonstrate that the grey mould fungus releases a large number of relatively non host-specific CDIPs and two toxins during infection, which collectively determine its necrogenic ability. We assume that the complexity and redundance of the phytotoxic secretome is correlated with the exceptionally wide host range and the ability of *B. cinerea* to successfully attack more than 1400 reported plant species (Elad Y *et al*. 2016). To gain a deeper insight into the mechanisms of host cell death induction, it is necessary to identify the receptors or targets of the CDIPs in order to study their mode of action.

## Materials and Methods

### Cultivation and transformation of *Botrytis*

*B. cinerea* strains were routinely cultured on agar containing malt extract (ME) medium (Müller *et al*. 2018). For growth tests, agar plates containing Gamborg minimal medium (GB5) containing 25 mM glucose were used and ME medium for sporulation tests. For these tests, 10 µl droplets containing 10^5^ conidia ml^-1^ were inoculated onto agar plates, and incubated at 20-22°C.

Classical transformation using knockout constructs with resistance markers was performed as described (Leisen *et al*. 2020; Müller *et al*. 2018). Protoplasts transformed with constructs containing CypR cassettes were selected in SH agar containing 0.3 µg ml^-1^ Cyp (Syngenta, Chorus® fungicide formulation). Non-transformed colonies appeared with a frequency of <10^−7^ per transformed protoplast. Resistant colonies were transferred after three days to plates containing GB5 agar with 25 mM glucose and 0.3 µg ml^-1^ Cyp. For generation of marker-free multiple knockout mutants, the transformation protocol of (Leisen *et al*. 2020) was modified as following: To 2× 10^7^ *B. cinerea* protoplasts suspended in 100 µl TMSC buffer, 10 µg pTEL-Fen and up to four RNPs, each consisting of pre-complexed 6 µg Cas9-Stu^2x^ and 2 µg sgRNA (two RNPs per gene) were added. The transformed protoplasts were mixed with liquified 200 ml SH agar adjusted to 39.5°C, and poured into ten 90 mm petri dishes. After three days of incubation at 20-22°C, small agarose pieces containing individual transformants were cut out with a scalpel and transferred to 5 cm plates containing selection-free 4x ME agar (4x ME: 4% malt extract, 1.6% glucose, 1.6% yeast extract, 1.5% agar, pH 5.5), to accelerate growth and sporulation of freshly generated transformants. After two days, Plates with no or very little growth were discarded. Hyphal tips of fast growing colonies were transferred to new 4x ME plates and allowed to grow for 5-6 days until sporulation. Conidia or sporulating mycelium were used for DNA isolation and PCR analysis to detect the desired editing events in each of the two genes and the absence of WT DNA in the deleted region. Transformants with deletions of one or both targeted genes were used for phenotypic characterization. Sometimes a single spore isolation was subsequently performed. The resulting culture was used for confirmation of the editing events, and proof of homokaryosis and absence of WT DNA using primers that amplified an internal sequence of the deleted regions.

### DNA manipulations

To generate a cyprodinil resistance (CypR) cassette, the *Bcpos5* (Bcin10g02880) coding sequence including two introns was amplified by PCR from genomic DNA of *B. cinerea* B05.10, by using primers CypR_ol_Ptrp_FW CypR_ol_TniaD_RV. The 3’-terminal primer CypR_ol_TniaD_RV was changed in sequence to generate the cypR-associated L412F substitution (Mosbach *et al*. 2017). The resulting fragment was flanked with fragments containing the *Aspergillus nidulans trpC* promoter (PtrpC) generated with primers Gib_pTEL_S_EcorV_PtrpC & PtrpC_ol_CypR_RV, and the *niaD* terminator of *B. cinerea* (TniaD) generated with primers TniaD_ol_CypR_FW & Gib_pTEL_S_EcorV_TniaD, both amplified from plasmid pTEL-Fen (Leisen *et al*. 2020). The CypR cassette was integrated into pTEL-Start linearized with EcoRV. To test its functionality as a resistance marker, the CypR cassette was attached to ca. 1 kb flanking regions of several target genes, using a modular cloning approach (see below). After transformation into *B. cinerea* protoplasts, selection was applied in SH agar with 0.3 µg ml^-1^ Cyp. Single spores of the transformants were transferred to Gamborg GB5 minimal medium with 25 mM glucose, supplemented with 0.3 µg ml^-1^ Cyp, for further cultivation and verification of the transformants.

Deletion constructs were generated with resistance cassettes for nourseothricin (natR/^N^) and cyprodinil (cypR/^C^) for *spl1*^*C*^, *xyn11A*^*N*^, *nep2*^*N*^, *ieb1* ^*C*^, *xyg1*^*C*^, *xyn11A*^*C*^, and *nep2*^*C*^. For this, 0.5 - 1 kb genomic regions flanking the coding sequences were amplified, spliced together with a resistance cassette (*trpC* promoter-resistance gene – *niaD* terminator) into a pBS-KS vector by Gibson assembly, and transformed into *E. coli*. Before transformation into *B. cinerea*, deletion constructs were released from the plasmids by restriction digestion.

For generation of a quadruple mutant, a *xyn11A*^*N*^ mutant was transformed with a *spl1*^*C*^ k.o. cassette to generate a *xyn11A*^*N*^ *spl1*^*C*^ double mutant. This mutant was cotransformed with two Cas9-sgRNA complexes targeting *nep1* and *nep2*, and *nep1*^*H*^ *nep2*^*F*^ k.o. cassettes with 60 bp homology flanks (amplified using pTEL-Fen or pTEL-Hyg as template) as repair templates. Transformants with resistance to FenR and HygR were tested for the knockout of *nep1* and *nep2*. The double mutant pg1pg2^R^, kindly provided by Jan van Kan (Wageningen University), was constructed transforming a hygR *pg1* mutant (Have *et al*. 1998) with a *pg2*^*N*^ knockout construct. The knockouts of *pg1* and *pg2* were confirmed by PCR. The primers used for synthesis of sgRNAs, construction of knockout constructs, and screening of transformants for correct knockouts and homokaryosis are shown in Table S3.

### Infection tests and secretome analyses

Infection tests were performed with attached leaves of *Phaseolus vulgaris* (genotype N9059), detached leaves of tomato (*Solanum lycopersicum*, cv. Marmande) and maize (*Zea mays*, cv. Golden Bantam), and to apple fruit (Malus domestica, cv. Golden Delicious). Leaf inoculations were performed as previously described (Schamber *et al*. 2010), using 20 µl droplets with 10^5^ conidia ml^-1^ in GB5 minimal medium (GB5: 3.05 g l-1 GB5, 10 mM KH_2_PO_4_, pH 5.5) with 25 mM glucose. To achieve maximal accuracy and comparability, two or three droplets each of WT and mutant conidial suspensions were applied on both sides of the midrib of one leaf or leaflet. Apple fruit was inoculated after wounding with a cork borer of 7 mm diameter along the equatorial line. Lesions were measured after 48 to 96 h by a caliper or image analysis using ImageJ software, and lesion areas calculated after subtraction of the inoculation area.

*On planta* secretomes were obtained from detached tomato leaves densely inoculated with 25 µl droplets containing 10^5^ conidia ml^-1^, and incubated at 20-22°C and 100% humidity in flat glass trays covered with saran wrap. After 48 h, droplets were collected, frozen at -80°C, thawed, centrifuged at 4°C for 60 min at 4000 g, sterile filtered and again frozen in aliquots at -80°C until further analysis. MS/MS-based proteomic analysis for confirmation of loss of CDIPs in the deletion mutants was performed as described (Müller *et al*. 2018). To determine CDI activity of WT and mutant secretomes, the secretomes (containing ca. 5-10 µg protein ml^-1^) were diluted with GB5 medium to concentrations of 1 or 2 µg protein ml^-1^, and ca. 20-50 µl each of the solutions were infiltrated into *Nicotiana benthamiana* or *Vicia faba* (cv. Fuego) leaves. The size of necrotic lesions in the infiltrated leaf area was recorded after two days. Heat treatments of the secretome were performed by incubation for 20 min at 95°C in a heating block. Size fractionation of the secretomes was done using ultrafiltration cartridges (Amicon Ultra-4, Merck Millipore Ltd, Tullagreen, Co., Cork, Ireland) with 10 kDa molecular weight cutoff. Before infiltration into leaves, the low molecular weight fraction was two-fold concentrated.

### Microscopic analysis of infection

To compare the early infection process of *B. cinerea* WT and multi-k.o. mutants microscopically, inoculations of detached *Phaseolus vulgaris* leaves and onion epidermal layers were performed. From onions, epidermal layers were removed from the concave side, fixed with tape onto glass coverslips and killed by incubation at 65°C for 30 min in a water bath. After thorough washing with water, samples were dried and inoculated with 20 µl droplets 5*10^4^ conidia ml^-1^ in 1 mM fructose, and incubated in a humid chamber for 24 h. For confocal microscopy, *Phaseolus vulgaris* leaves were inoculated with 1 µl of 10^5^ conidia ml^-1^ in GB5 medium with 25 mM glucose. After 24 h, developing lesions were stained with 10µg ml^-1^ calcofluor white (fluorescence brightener 28, Sigma) for 5 min and thoroughly washed subsequently. Images were acquired using a Zeiss LSM880 AxioObserver confocal laser scanning microscope equipped with a Zeiss EC Plan-Neofluar 5x/0.16 M27 objective (DFG, INST 248/254-1). Fluorescent signals of calcofluor white (excitation/emission 405 nm/410-523-571 nm) and chlorophyll (excitation/emission 633nm/638-721 nm) were processed using the Zeiss software ZEN 2.3 or ImageJ (https://imagej.nih.gov/ij/).

### Statistical analyses

Statistical analyses were carried out with the GraphPad Prism software. For comparison of radial growth, means of three experiments, with three replicates each, were analysed by one way ANOVA followed by Dunnett’s multiple comparisons post-hoc test. For infection assays, two or three pairwise inoculations of WT and mutant were performed on the same leaf or fruit (Fig. 2). Relative necrotic areas (% of WT) were calculated for each leaf/fruit, based on technical replicates of WT vs. mutant pairs. Values from at least three inoculation dates with at least three leaves/fruits each were analyzed by one-sample t test. Comparison of relative necrotic areas between different deletion mutants was carried out by one-way ANOVA followed by Tukey’s multiple comparison test. Box limits of box plots represent 25th percentile and 75th percentile, horizontal line represents median, and whiskers display minimum to maximum values. Evaluation of MS/MS proteomics data for significant deviations in the abundance of detected proteins in WT and 10x, 11x, 12xbb and 12xpg mutants was performed using the Perseus bioinformatics platform (Tyanova *et al*. 2016).

## Acknowledgments

We are grateful to Jan van Kan for providing us the *pg1 pg2* mutant, and to Sophie Eisele and Sarah Gabelmann for help in characterizing the mutants. We also thank Olivia Reichle, Jonas Müller, Sabrina Kaiser and Jacqueline Hackh who helped with the acquisition and evaluation of the secretome data and microscopic pictures. Special thanks to Andrew Foster, who suggested the use double RNPs without repair templates for efficient gene knockouts, and to Felix Willmund and A. Sharon for helpful comments on the manuscript. This work was supported by the BioComp Research Initiative of Rhineland-Palatinate, Germany.

## Supplemental Figures and Tables

**Figure S1:**
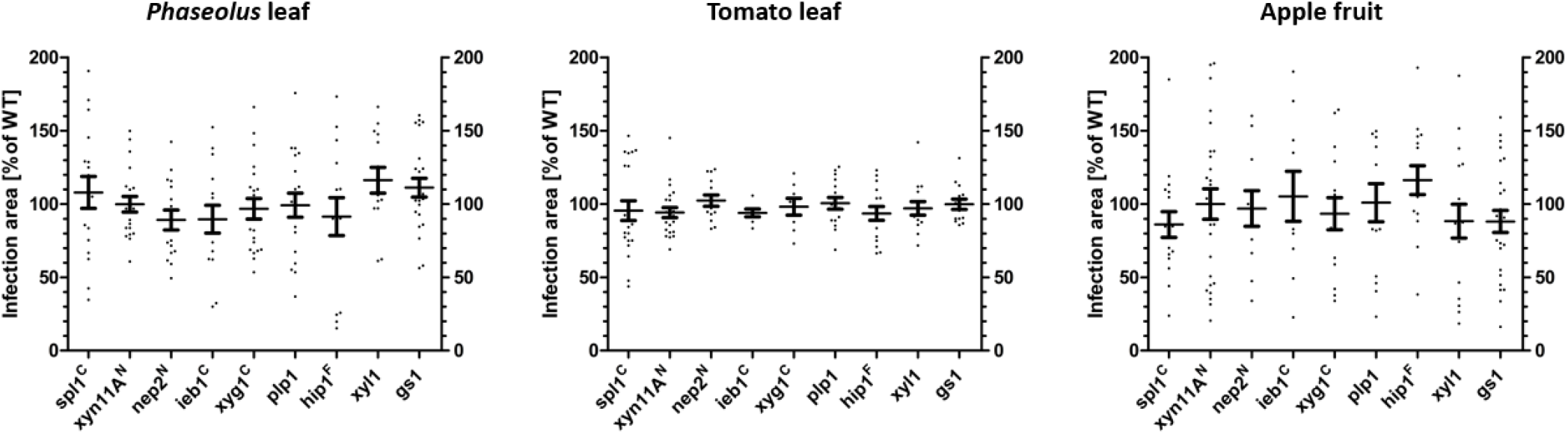
Infection tests with single CDIP mutants. The p values by one-sample t test to a hypothetical value of 100% (WT) did not show significant diferences for any of the tested mutants.

**Figure S2:**
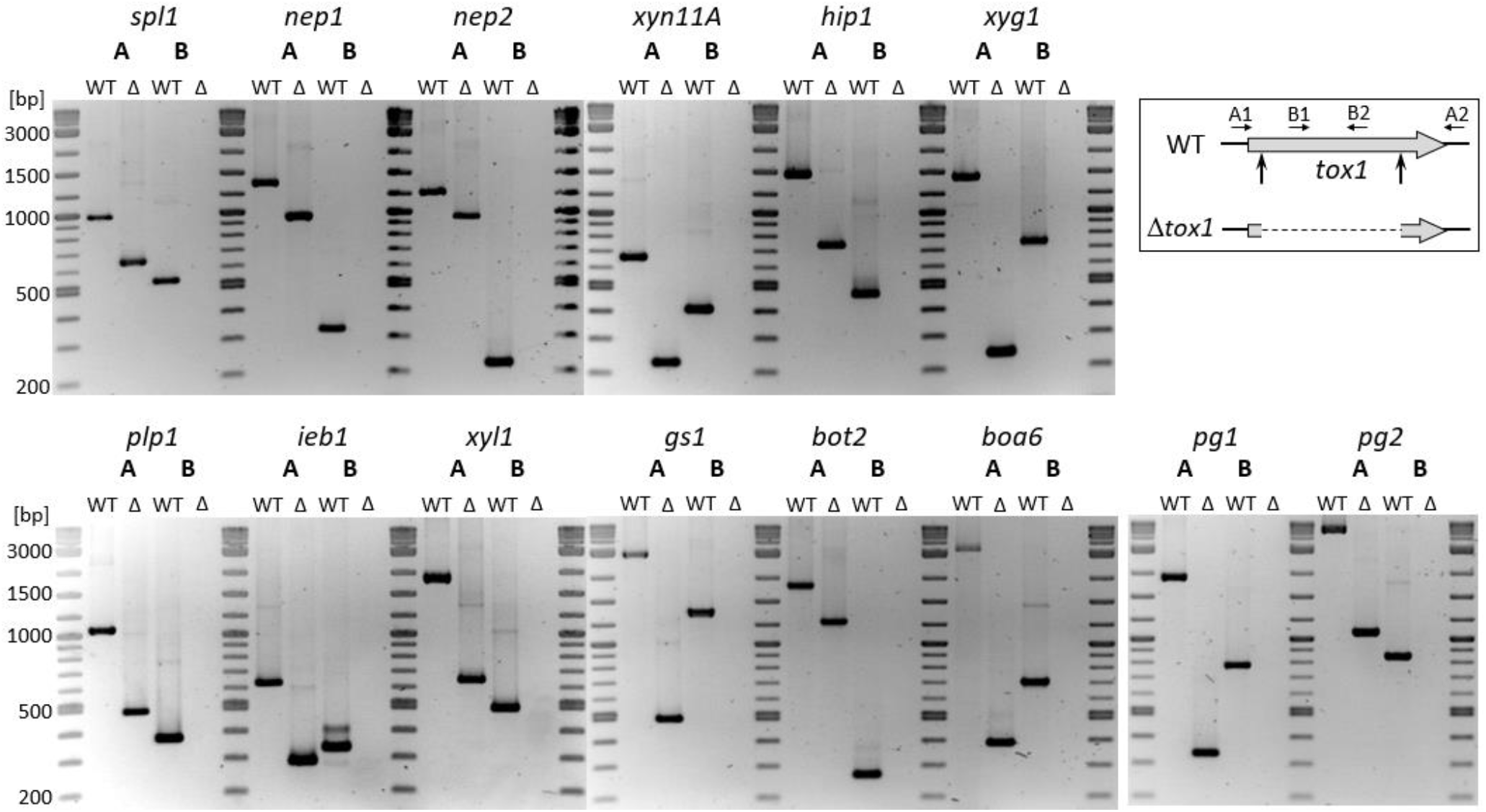
PCR-based confirmation of gene deletions in the 12xbb mutant. Position of primer pairs used are indicated in the sketch. A: PCR with primers flanking the deleted region B: PCR with primers amplifying an internal part of the deleted region. Missing PCR products in reactions B confirm homokaryosis of deletion mutants. The size of the deletions was determined by sequencing (Table S2). Primers used are shown in Table S4. Mapping of *pg1* and *pg2* deletions was done with the 12xpg mutant.

**Figure S3:**
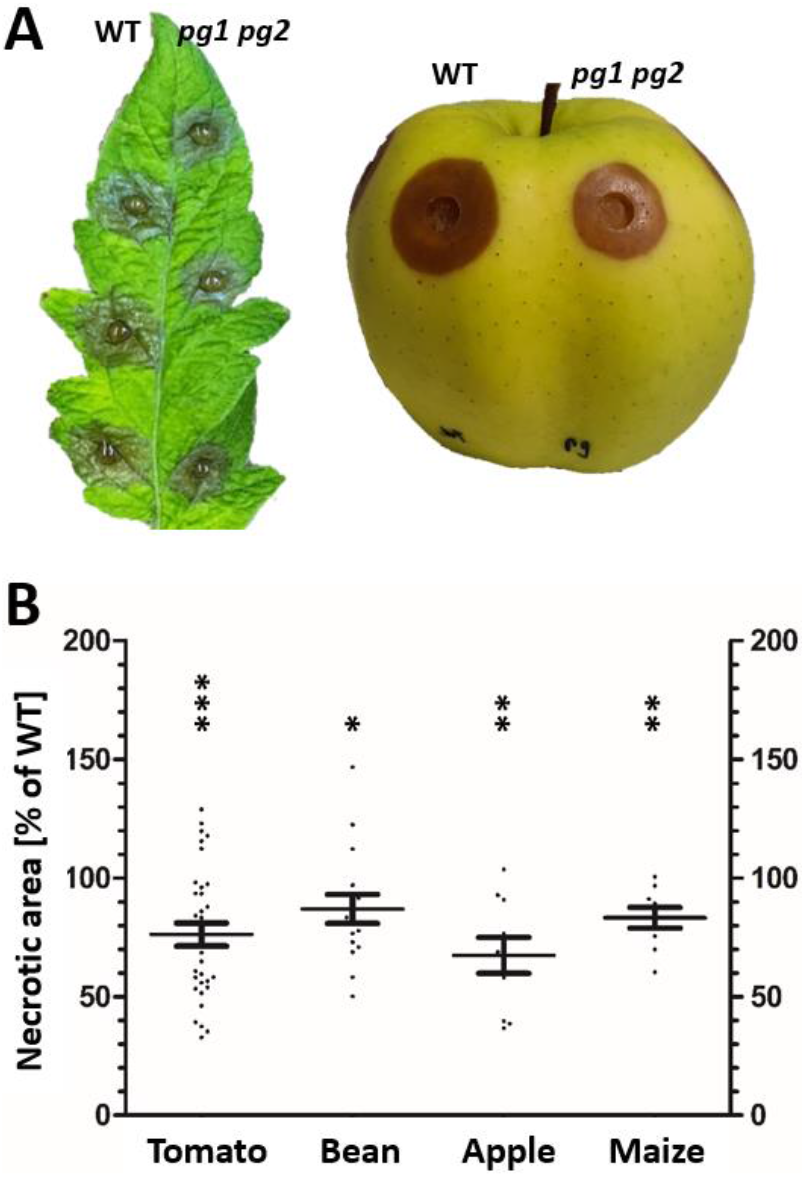
Characterization of a *pg1 pg2* double mutant (provided by J. van Kan). A:: Infected tomato leaf (72 h) and apple fruit (96 h). B: Results of infection tests on different plant tissues (cf. Fig. 1). The p values by one-sample t test to a hypothetical value of 100% (WT) are indicated. * p value < 0.05; ** p value < 0.01; *** p value < 0.001.

**Figure S4:**
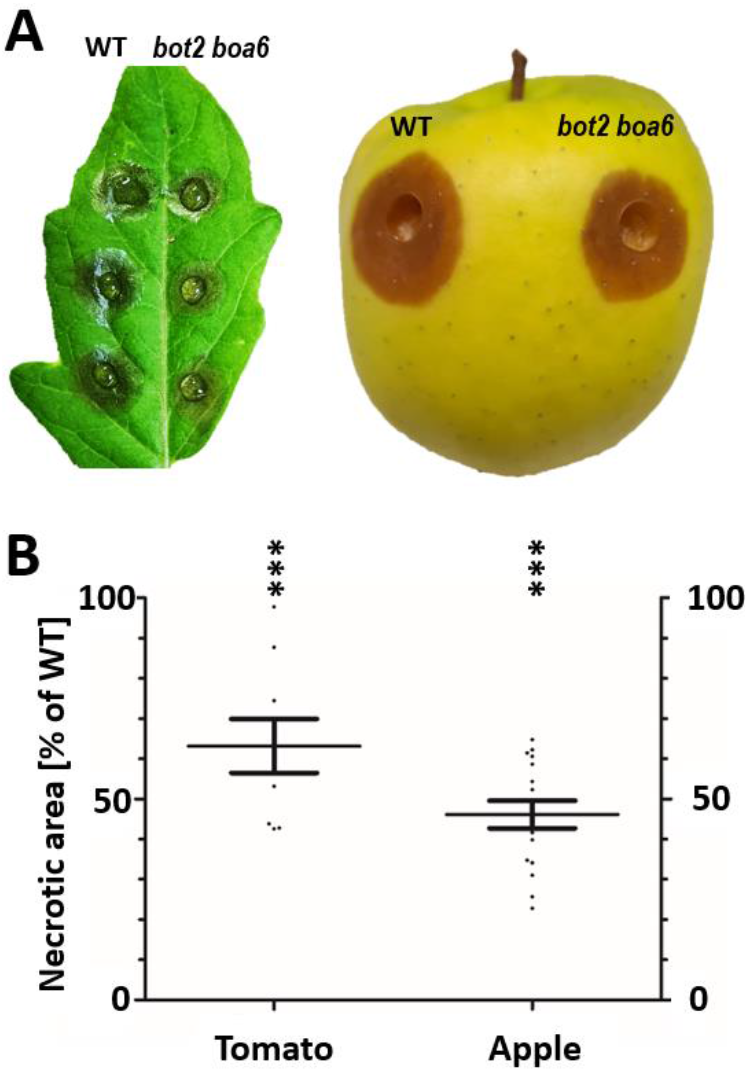
Lesion formation of the *B. cinerea bot2 boa6* mutant (Leisen *et al*. 2020) on tomato leaf (48 h) and apple fruit (96 h), compared to WT. The p values by one-sample t test to a hypothetical value of 100% (WT) are indicated *** p value < 0.001.

**Figure S5:**
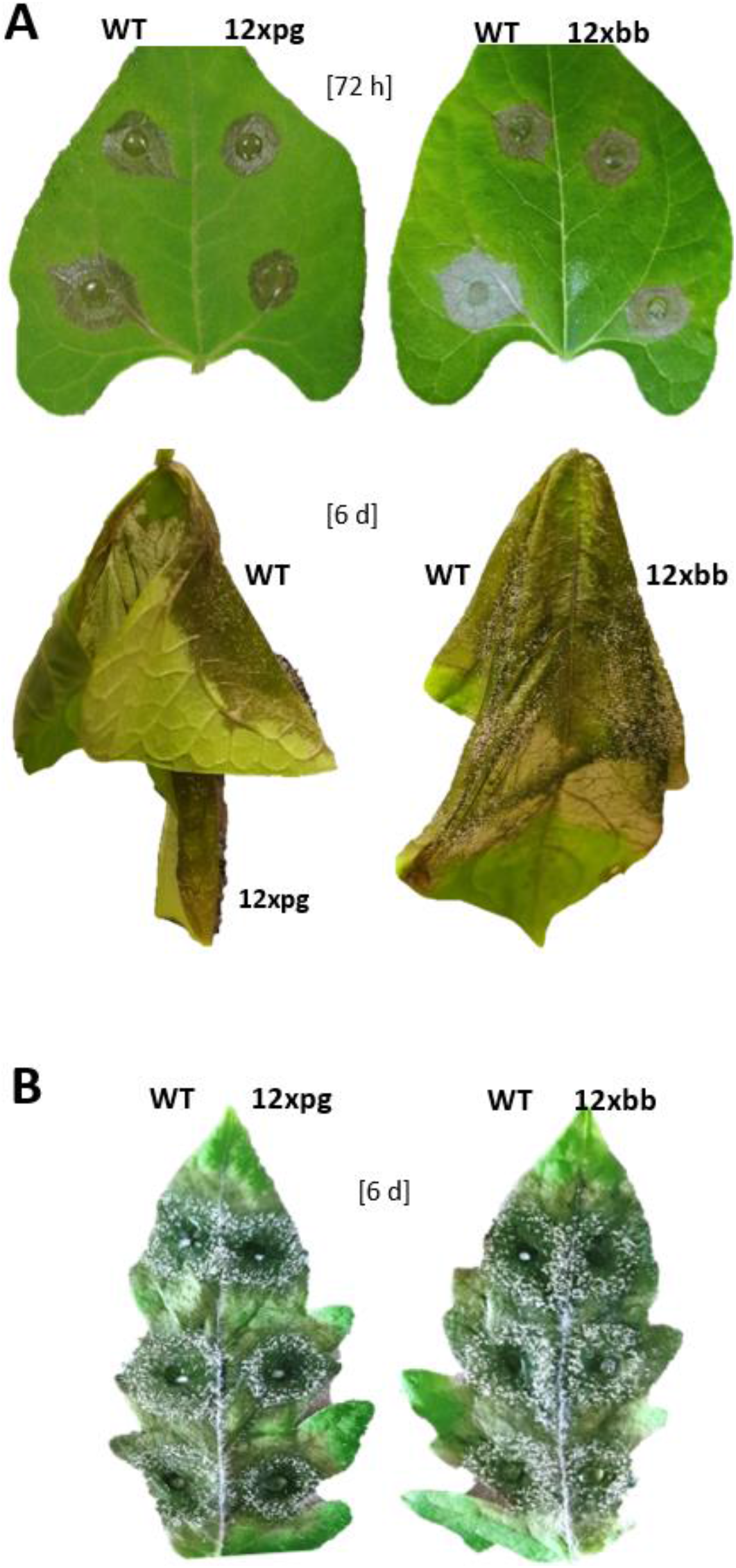
Lesion formation and sporulation of *B. cinerea* WT, 12xpg and 12xbb mutants on attached. *Phaseolus* bean leaves (A: 3 days; B: 6 days), and on detached tomato leaves (6 days).

**Table S1:** Mapping of the deletions in the 12xbb and 12xpg (*pg1* and *pg2* only) mutants.

| Gene | Accession | Protein size (aa) | Deletion mapping |  |  |  |
| --- | --- | --- | --- | --- | --- | --- |
|  |  |  | Predicted | Obtained | Size (change) | Deleted codons |
| *spl1* | Bcin03g00500 | 137 | 3:185840-186170 | as expected | 331 bp | 24-end* |
| *nep1* | Bcin06g06720 | 246 | 6:2351724-2352121 | as expected | 398 bp | 34-end* |
| *nep2* | Bcin02g07770 | 244 | 2:2793732-2794002 | as expected | 271 bp | 80-133 |
| *xyn11A* | Bcin03g00480 | 227 | 3:174482-174915 | as expected | 434 bp | 64-end* |
| *hip1* | Bcin14g01200 | 151 | 14:538686-539428 | 14:538687-539428 | 742 bp (-1) | all |
| *xyg1* | Bcin03g03630 | 248 | 3:1224165-1225358 | 3:1224166-1225358 | 1193 bp (-1) | all |
| *plp1* | Bcin10g01020 | 147 | 10:417795-418359 | 10:417796-418359 | 564 bp (-1) | all |
| *ieb1* | Bcin15g00100 | 187 | 15:78364-78704 | as expected | 341 bp | 20-115 |
| *xyl1* | Bcin09g01800 | 329 | 9:672899-674137 | as expected | 1239 bp | all |
| *gs1* | Bcin04g04190 | 645 | 4:1491835-1494178 | as expected | 2344 bp | all |
| *bot2* | Bcin12g06390 | 399 | 12:2224281-2224868 | as expected | 588 bp | 157-end* |
| *boa6* | Bcin01g00060 | 2460 | 1:18020-20645 | as expected | 2626 bp | 661-end* |
| *pg1* | Bcin14g00850 | 382 | 14:371069-372695 | 14:371071-372695 | 1625 bp (-2) | all |
| *pg2* | Bcin14g00610 | 374 | 14:247707 to 249005 | 14:245745 to 250019 | 1299 / 4275** | all |

Deletions were predicted based on the expected cleavage sites of each pair of Cas9-RNP. \* Deletion of codons due to frameshift. \*\*The 4275 bp (instead of 1299 bp) deletion extended into Bcin14g00600 encoding a sclerotia-specifically expressed polyketide synthetase. Since this gene is not expressed *in planta*, it is not expected to contribute to infection.

**Table S2:** MS/MS-detection of CDIPs in the secretomes of *B. cinerea* WT and multiple k.o. mutants.

|  | n |  | Spl1* | Nep2 | Xyn11A | Hip1 | XYG1 | IEB1 | Gs1 | PG1* | PG2* |
| --- | --- | --- | --- | --- | --- | --- | --- | --- | --- | --- | --- |
| WT | 6 | mean | 560.332 | 41.818 | 337.044 | 48.416 | 595.662 | 509.355 | 406.258 | 697.042 | 1.184.440 |
|  |  | st.dev. | 409.251 | 21.637 | 377.637 | 81.950 | 393.979 | 351.129 | 282.887 | 856.712 | 1..157307 |
| 4x<sup>R</sup> | 5 | mean | 0 | 0 | 0 | 8.355 | 609.594 | 564.152 | 475.172 | 937.821 | 1.213.902 |
|  |  | st.dev. | 0 | 0 | 0 | 16.710 | 489.109 | 553.043 | 299.148 | 1.306.651 | 1.017.928 |
| pg1 pg2 | 4 | mean | 377.472 | 25.224 | 273.638 | 13.480 | 931.655 | 415.655 | 436.435 | 274 | 250 |
|  |  | st.dev. | 420.371 | 16.477 | 226.923 | 23.348 | 825.703 | 190.870 | 306.249 | 474 | 432 |
| WT | 5 | mean | 4.292.542 | 712.180 | 9.987.220 | 269.486 | 2.946.020 | 4.583.840 | 3.756.580 | 7.239.160 | 5.246.700 |
|  |  | st.dev. | 2.427.053 | 177.747 | 1.684.412 | 330.555 | 935.540 | 1.328.030 | 498.726 | 1.554.149 | 2.604.637 |
| 10x | 4 | mean | 1.771 | 0 | 0 | 0 | 0 | 0 | 0 | 880.6725 | 2.772.025 |
|  |  | st.dev. | 3.067 | 0 | 0 | 0 | 0 | 0 | 0 | 2.207.591 | 58.036 |
| 11x | 3 | mean | 32.047 | 0 | 0 | 0 | 0 | 0 | 0 | 16.757 | 12.606.067 |
|  |  | st.dev. | 24.337 | 0 | 0 | 0 | 0 | 0 | 0 | 23.698 | 2.980.671 |
| 12xpg | 3 | mean | 3.545 | 0 | 0 | 0 | 0 | 0 | 0 | 27.588 | 19.550 |
|  |  | st.dev. | 5.013 | 0 | 0 | 0 | 0 | 0 | 0 | 12.016 | 27.648 |
| 12xbb | 3 | mean | 46.996 | 0 | 0 | 0 | 0 | 0 | 0 | 7.956.433 | 9.498.333 |
|  |  | st.dev. | 23.050 | 0 | 0 | 0 | 0 | 0 | 0 | 1.672.172 | 6.315.201 |

Mean LQF intensities are shown. Proteomic analysis was performed in two series of experiments, the first series including WT, 4x<sup>R</sup> and *pg1 pg2* mutants, and the second series WT, 10x, 11x, 12xpg and 12xbb mutants. n: Number of replicates. Values of proteins that have been deleted in the analysed mutants are shown with red background. \*The low intensity values observed for Spl1, PG1 and PG2 are probably due to carry-over of some peptides from WT samples during MS/MS analysis.

**Table S3:**
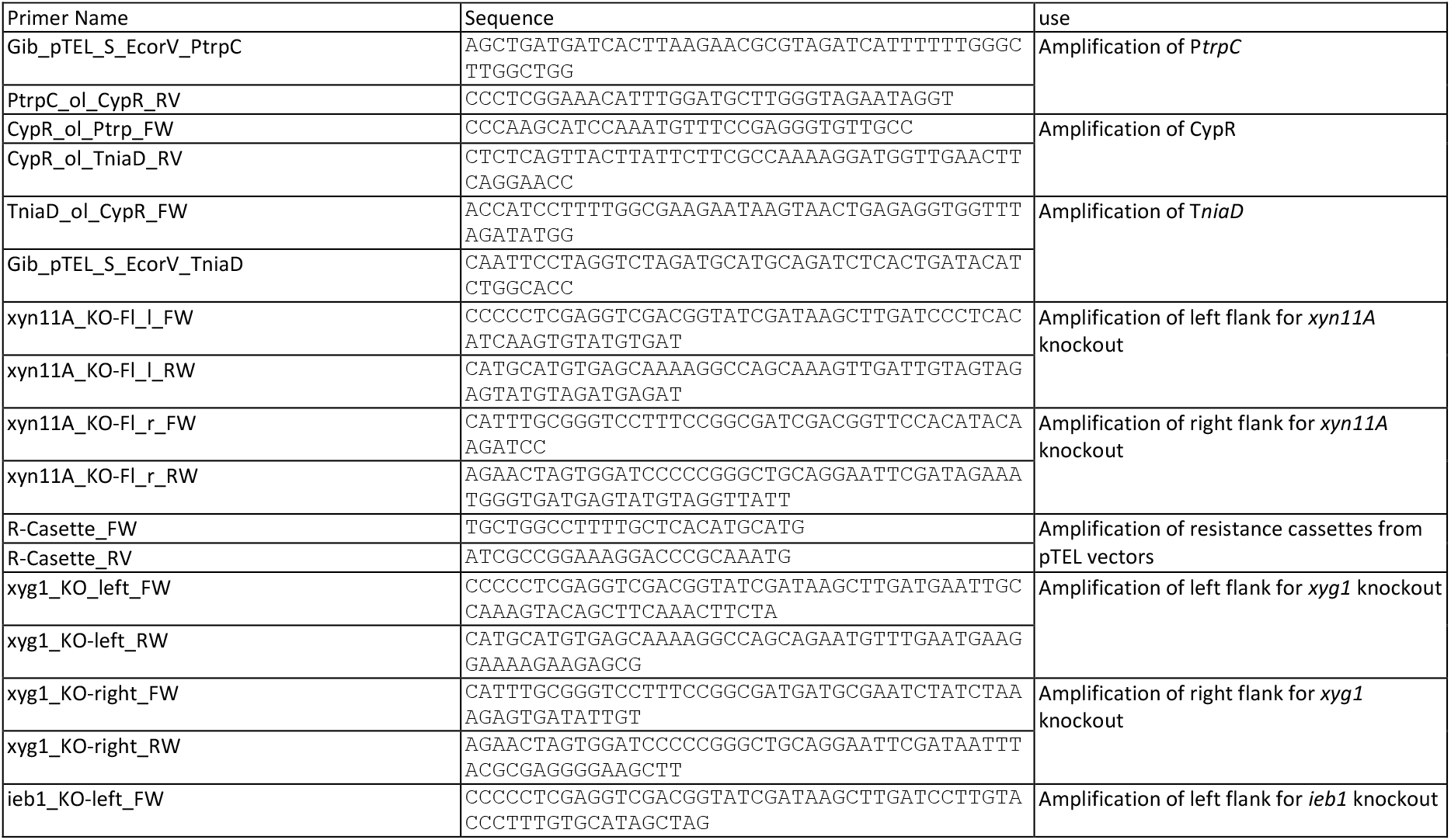
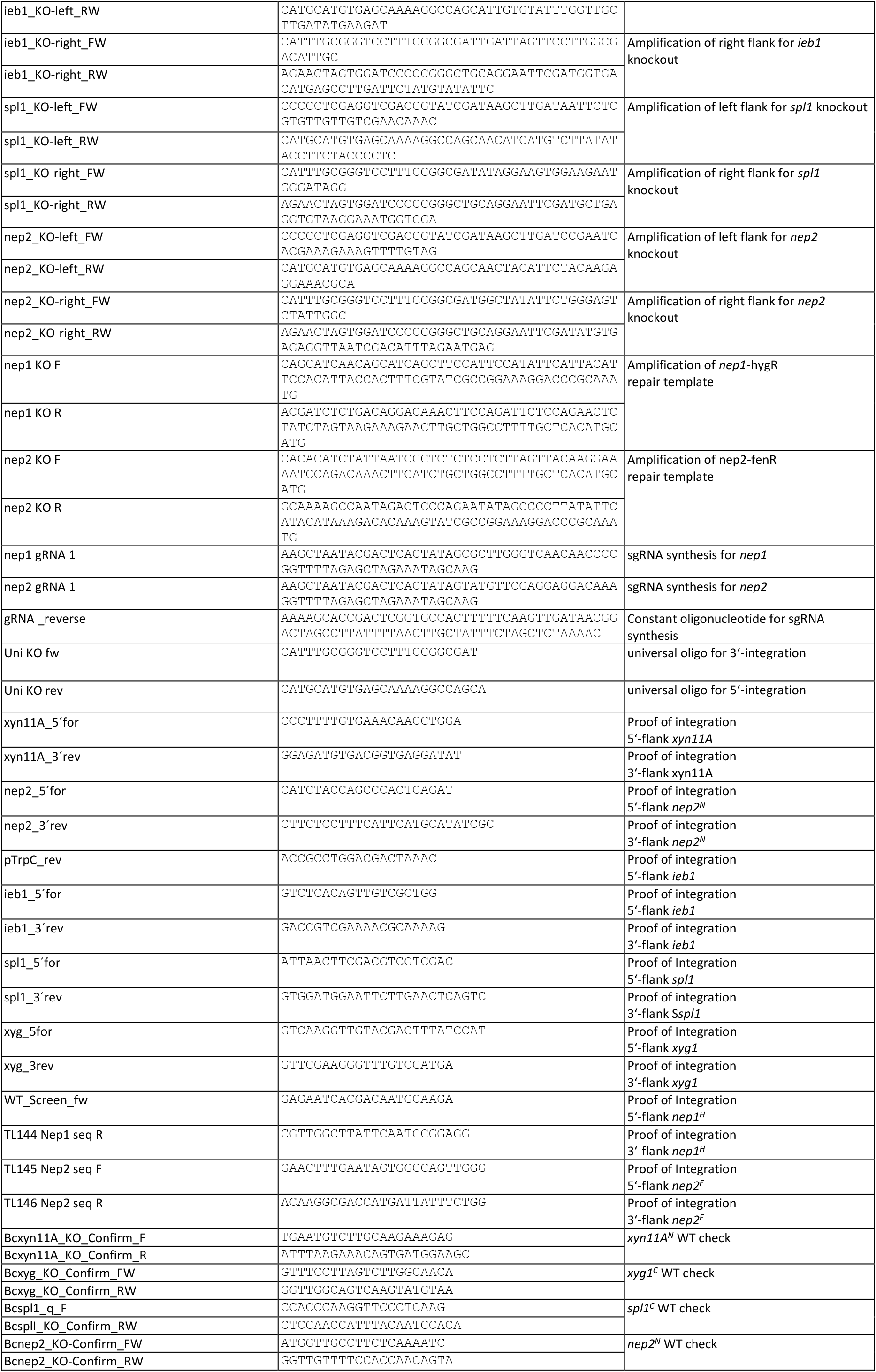
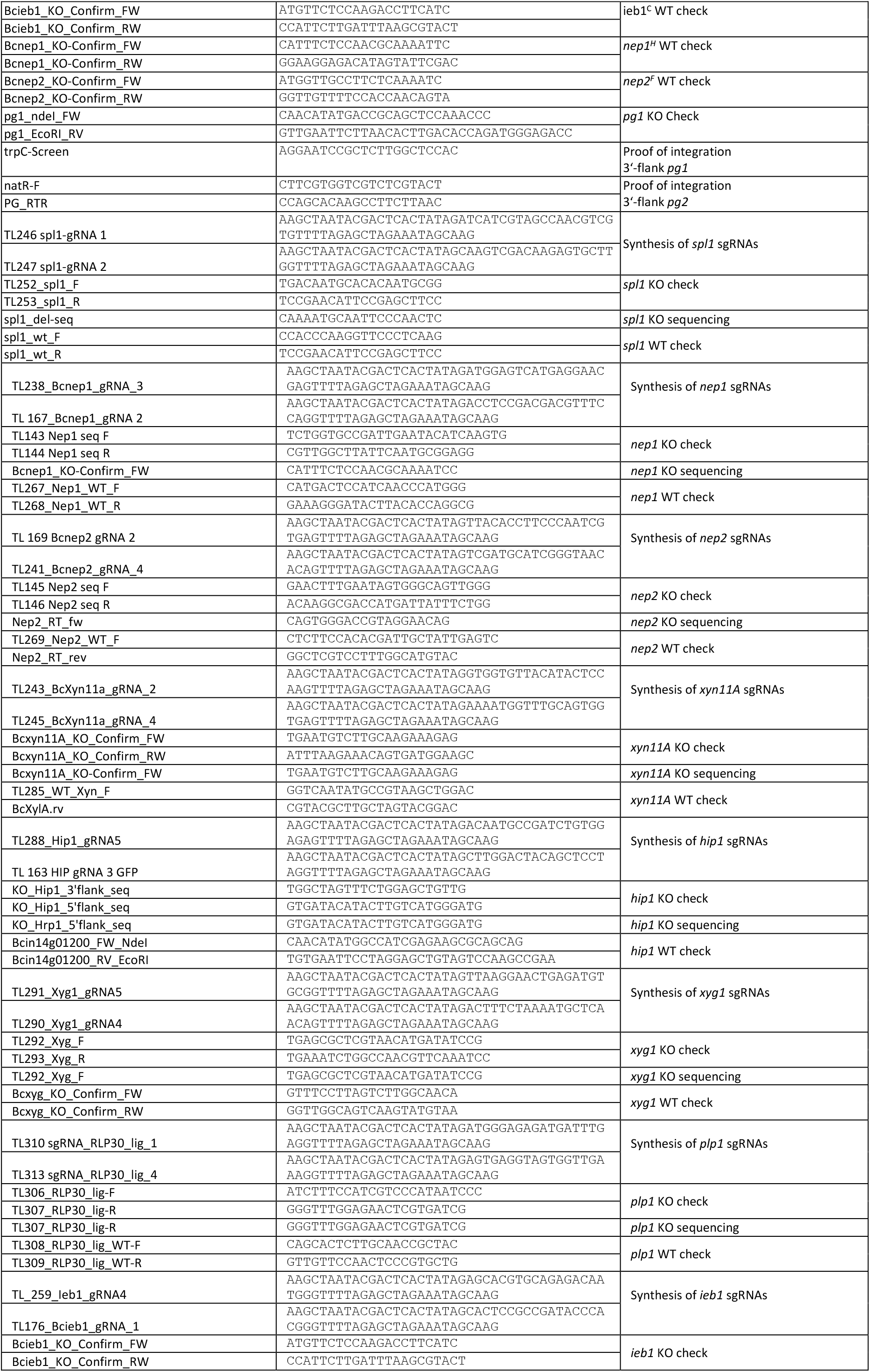
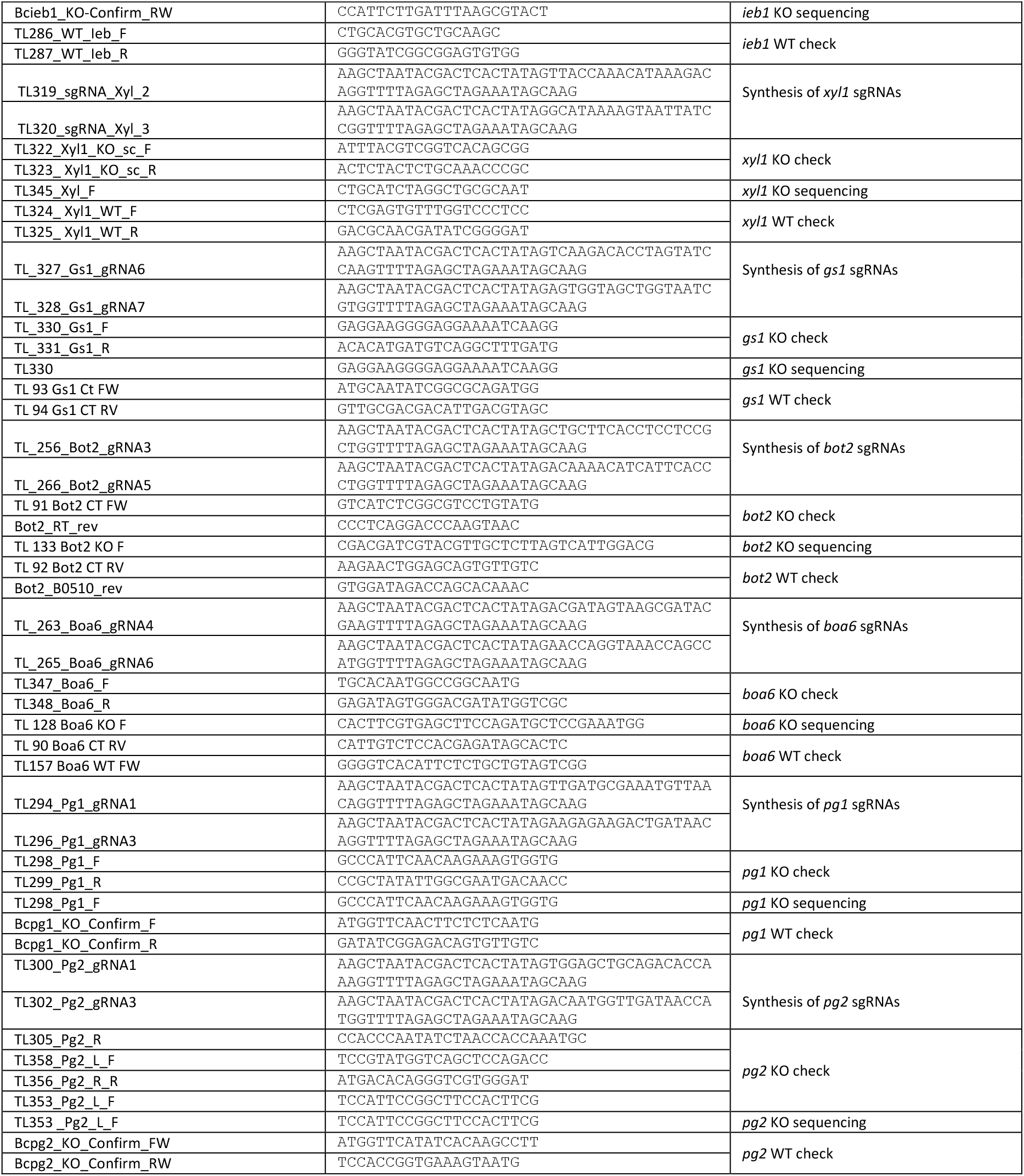
Oligonucleotides used.

